# OncodriveCLUSTL: a sequence-based clustering method to identify cancer drivers

**DOI:** 10.1101/500132

**Authors:** Claudia Arnedo-Pac, Loris Mularoni, Ferran Muiños, Abel Gonzalez-Perez, Nuria Lopez-Bigas

## Abstract

**Summary:** The identification of the genomic alterations driving tumorigenesis is one of the main goals in oncogenomics research. Given the evolutionary principles of cancer development, computational methods that detect signals of positive selection in the pattern of tumor mutations have been effectively applied in the search for cancer genes. One of these signals is the abnormal clustering of mutations, which has been shown to be complementary to other signals in the detection of driver genes. We have developed OncodriveCLUSTL, a new sequence-based clustering algorithm to detect significant clustering signals across genomic regions. OncodriveCLUSTL is based on a local background model derived from the simulation of mutations accounting for the composition of tri- or penta-nucleotide context substitutions observed in the cohort under study. Our method is able to identify known clusters and *bona-fide* cancer drivers across cohorts of tumor whole-exomes, outperforming the existing OncodriveCLUST algorithm and complementing other methods based on different signals of positive selection. We show that OncodriveCLUSTL may be applied to the analysis of non-coding genomic elements and non-human mutations data.

**Availability and implementation:** OncodriveCLUSTL is available as an installable Python 3.5 package. The source code and running examples are freely available at https://bitbucket.org/bbglab/oncodriveclustl under GNU Affero General Public License.

## 1 Introduction

The identification of the alterations driving tumorigenesis is a major goal of cancer research. The knowledge of the molecular mechanisms underlying tumorigenesis is a necessary step for the implementation of precision cancer medicine. Cancer development is an evolutionary process; thus the detection of signals of positive selection in the somatic mutational pattern of genes has been exploited to identify drivers across tumor cohorts. Specifically, the non-random spatial accumulation, clustering, of mutations along the protein sequence has been used to identify cancer drivers and provide clues of oncogenic mechanisms (Tamborero *et al*., 2013a; Chang *et al*., 2016; Tokheim *et al*., 2016). This signal is complementary to other (such as recurrence) and thus, their combination can produce more comprehensive lists of driver genes (Tamborero *et al*., 2013b; Porta-Pardo *et al*., 2017).

Since the rate of generation of mutations across the genome is highly variable (Stamatoyannopoulos *et al*., 2009; Schuster-Böckler and Lehner 2012; Alexandrov *et al*., 2013; Lawrence *et al*., 2013; Polak, *et al*., 2015), clustering-based methods face the challenge to construct an accurate background model of the distribution of mutations to correctly assess the significance of observed clusters. Ideally, such model would include all the genomic position-dependent covariates of the mutation rate. Alternatively, one can locally simulate the same number of mutations as observed in the region following the probabilities of k-nucleotide context-dependent substitutions and assess weather the distribution of mutations along the region follows the expected (Mularoni *et al*., 2016). This background model is not affected by large-scale covariates of the mutation rate (e.g., replication timing or chromatin state) and may thus be applied to any region of the genome of any species.

Here we introduce OncodriveCLUSTL, a new linear clustering algorithm to detect genomic regions and elements with significant clustering signals based on a local background model derived from a cohort’s observed tri- or penta-nucleotide substitutions frequency. OncodriveCLUSTL identifies known mutation clusters and driver genes across TCGA cohorts, overperforming the existing OncodriveCLUST (Tamborero *et al*., 2013a), and complementing methods based on different signals of positive selection. We show that OncodriveCLUSTL identifies mutation clusters in promoter regions and mouse genes.

## 2 Implementation and availability

OncodriveCLUSTL is an unsupervised clustering algorithm implemented in Python 3.5. It analyzes somatic mutations that have been observed in genomic elements (GEs) across a cohort of samples (Fig.1A-1). Mutations in each GE are smoothed along its sequence using a Tukey based kernel density function and clusters are identified (Fig.1A-2,3), with scores based on the number and distribution of their mutations. Cluster scores are summed up to produce a GE clustering score. The significance of the observed clusters and GEs is assessed through *n* iterations of randomly distributed each GE observed mutation within a window of nucleotides centered at its position, following the frequency of cohort tri- or penta-nucleotide changes (Fig.1A-4,5; Supp. Methods for further details). P-values are adjusted using the Benjamini-Hochberg method at 1% false-discovery rate (FDR). OncodriveCLUSTL source code and examples are freely available at https://bitbucket.org/bbglab/oncodriveclustl. A web version of OncodriveCLUSTL can be run at https://bbglab.irbbarcelona.org/oncodriveclustl.

**Fig. 1.**
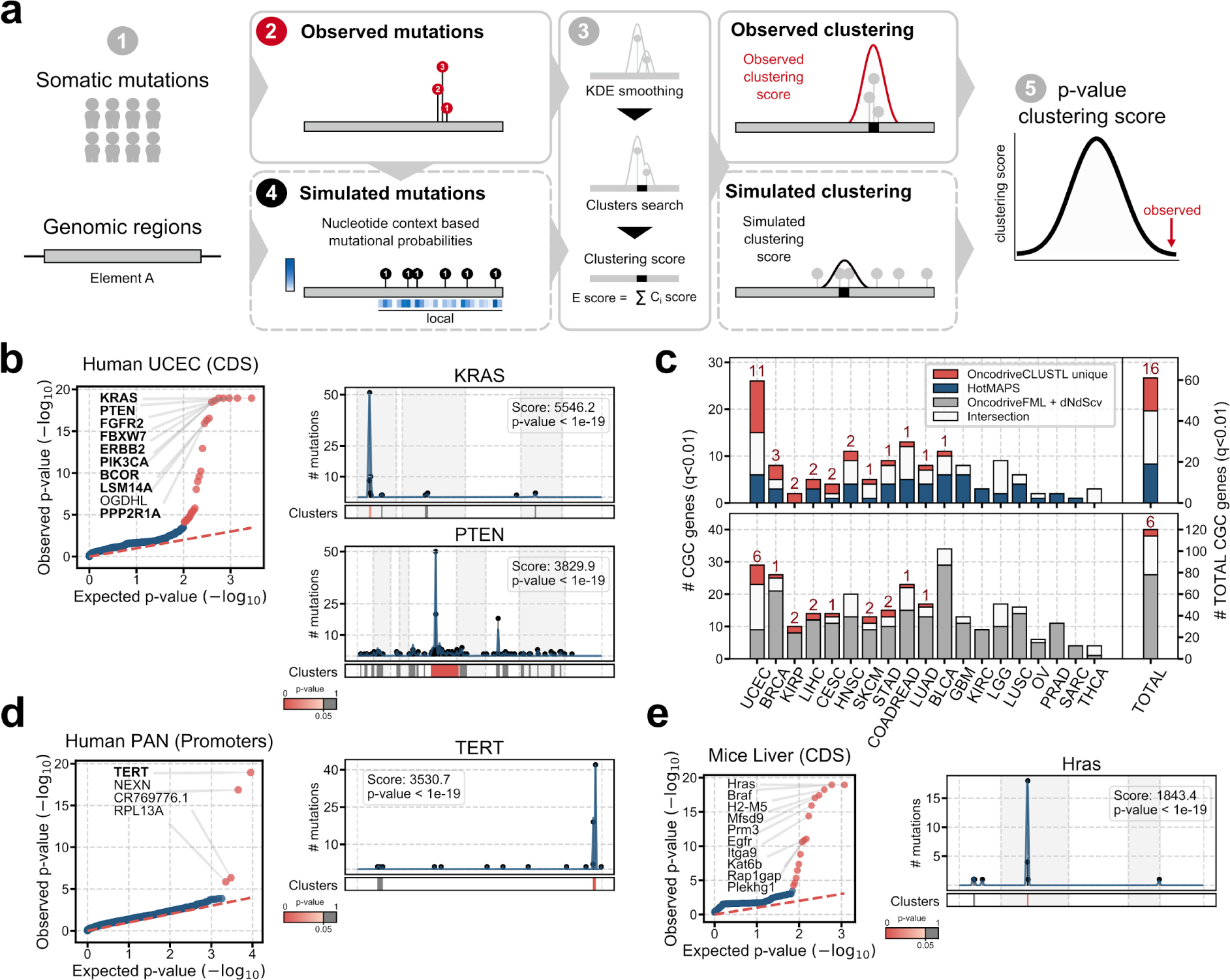
OncodriveCLUSTL algorithm and results. Overview of OncodriveCLUSTL (a). OncodriveCLUSTL detects well known cancer genes (b) and complements methods based on different signals of positive selection (c). OncodriveCLUSTL can be successfully applied to mutations in promoter regions (d) and mice genes (e).

## 3 Performance

### Mutations in human protein-coding genes across 19 TCGA cohorts

(Ellrott *et al*., 2018). OncodriveCLUSTL detects well-known cancer genes (CGC; Sondka *et al.*, 2018) with clusters of different sizes (Fig.1B; Supp.Fig.3-7). It outperforms the previously developed OncodriveCLUST (Tamborero *et al*., 2013a) in all cohorts (Supp.Fig.8), demonstrating that the improved clustering detection method and background model fine-tune the detection of drivers. It also exhibits similar performance than the 3D protein-clustering method HotMAPS (Tokheim *et al*., 2016) (Supp.Fig.8) detecting bona-fide drivers missed by the 3D analysis (Fig.1C). The results of OncodriveCLUSTL complement those of methods based on different signals of positive selection (OncodriveFML, Mularoni *et al*., 2016; dNdScv, Martincorena *et al*., 2017) (Fig.1C), thus highlighting the relevance of combining methods exploiting different signals for comprehensiveness.

### Mutations in promoters across a cohort of tumor whole-genomes

(Fredriksson *et al*., 2014). Consistent with the study describing the dataset, OncodriveCLUSTL found a significant cluster in the TERT promoter (Fig.1D), the mutations of which result in the upregulation of TERT (Supp.Fig.9). Significant clustering is also detected in few other promoters, which need to be carefully vetted to nominate them as cancer drivers, as we and others have shown that some local mutational processes may also lead to mutation clustering (Sabarinathan *et al*., 2016; Zou *et al*., 2017).

### Mutations in C3H mice genes in chemically-induced hepatocarcinomas (Connor *et al*., 2018)

As described by the authors of the dataset, OncodriveCLUSTL identified significant clustering in Hras, Braf and Egfr (Fig.1E).

## 4 Conclusions

OncodriveCLUSTL is a new method to identify sequence-based clustering signals across the genome. It operates with satisfactory sensitivity and specificity, outperforming the existing OncodriveCLUST and complementing other methods of driver detection in coding sequences. It may also be successfully applied to the detection of mutational clustering in non-coding regions and in non-human data.

## Acknowledgements

The results shown here are in part based upon data generated by the TCGA Research Network: https://cancergenome.nih.gov/.

## Funding

This work was supported by funding from the Spanish Ministry of Economy and Competitiveness [SAF2015-66084-R, MINECO/FEDER, UE] and by the European Research Council [Consolidator Grant 68239]. IRB Barcelona is the recipient of a Severo Ochoa Centre of Excellence Award from the Spanish Ministry of Economy and Competitiveness (MINECO; Government of Spain) and is supported by CERCA (Generalitat de Catalunya). A.G.-P. is supported by a Ramón y Cajal contract from the Spanish Ministry of Economy and Competitiveness [RYC-2013-1455]. C.A.-P. is supported by “La Caixa” Foundation [B004697].

## References

Alexandrov, L.B. et al. (2013) Signatures of mutational processes in human cancer. Nature, 500, 415–421.

Chang, M.T. et al. (2016) Identifying recurrent mutations in cancer reveals widespread lineage diversity and mutational specificity. Nat. Biotechnol., 34, 155–163.

Connor, F. et al. (2018) Mutational landscape of a chemically-induced mouse model of liver cancer. J. Hepatol., 69, 840–850.

Fredriksson, N.J. et al. (2014) Systematic analysis of noncoding somatic mutations and gene expression alterations across 14 tumor types. Nat. Genet., 46, 1258–1263.

Lawrence, M.S. et al. (2013) Mutational heterogeneity in cancer and the search for new cancer-associated genes. Nature, 499, 214–218.

Martincorena, I. et al. (2017) Universal Patterns of Selection in Cancer and Somatic Tissues. Cell, 171, 1029–1041.e21.

Mularoni, L. et al. (2016) OncodriveFML: A general framework to identify coding and non-coding regions with cancer driver mutations. Genome Biol., 17, 128.

Porta-Pardo, E. et al. (2017) Comparison of algorithms for the detection of cancer drivers at subgene resolution. Nat. Methods, 14, 782–788.

Sabarinathan, R. et al. (2016) Nucleotide excision repair is impaired by binding of transcription factors to DNA. Nature, 532, 264–267.

Sondka, Z. et al. (2018) The COSMIC Cancer Gene Census: describing genetic dysfunction across all human cancers. Nat. Rev. Cancer, 18, 696–705.

Stamatoyannopoulos, J.A. et al. (2009) Human mutation rate associated with DNA replication timing. Nat. Genet., 41, 393–395.

Tamborero, D. et al. (2013) OncodriveCLUST: Exploiting the positional clustering of somatic mutations to identify cancer genes. Bioinformatics, 29, 2238–2244.

Tamborero, D. et al. (2013) Comprehensive identification of mutational cancer driver genes across 12 tumor types. Sci. Rep., 3, 2650.

Tokheim, C. et al. (2016) Exome-scale discovery of hotspot mutation regions in human cancer using 3D protein structure. Cancer Res., 76, 3719–3731.

Zou, X. et al. (2017) Short inverted repeats contribute to localized mutability in human somatic cells. Nucleic Acids Res., 45, 11213–11221.

