## Supplementary material for "OncodriveCLUSTL: a sequence-based clustering method to identify cancer drivers": Methods and Supplementary Material

[1]Institute for Research in Biomedicine (IRB Barcelona), The Barcelona Institute of Science and Technology (BIST), Barcelona, Spain.
[2]Research Program on Biomedical Informatics, Universitat Pompeu Fabra, Barcelona, Spain.
[3]Institució Catalana de Recerca i Estudis Avançats (ICREA), Passeig Lluís Companys 23, 08010 Barcelona, Spain.
[#]To whom correspondence should be addressed.

### Index

### 1 Supplementary methods

#### Tumor mutation datasets

We obtained the mutations identified in primary tumors from three different datasets: 19 whole exome sequencing (WXS) cohorts of The Cancer Genome Atlas (TCGA) project (Ellrott *et al.*, 2018) (downloaded on December 29th, 2016), 14 whole genome sequencing cohorts from TCGA WGS-505 project (Fredriksson *et al.*, 2014) (downloaded on March 31st, 2015), and a WXS dataset (European Nucleotide Archive ERZ537501) of chemically-induced C3H mice liver tumors (Connor *et al.*, 2018) (downloaded on October 11th, 2018). TCGA WXS cohorts consist of 8,263 samples (1,382,259 substitutions) from 19 different cancer types, each of them composed by a minimum of 200 samples. TCGA WGS-505 consist of 505 samples (12,423,016 substitutions) from 14 different cancer types. A whole-genome Pancancer cohort was obtained by merging all cohorts in the TCGA WGS-505 dataset. Hypermutated tumors were removed from the datasets. To define hypermutated samples for a given cohort, we calculated the distribution of the number of alterations across samples in the cohort. Those samples bearing a minimum number of 1,000 alterations exceeding 1.5 times the interquartile range above the 75th percentile were considered hypermutated. Original mouse WXS dataset consist of substitutions identified across 78 liver tumors of C3H mice generated spontaneously or by exposure to diethylnitrosamine (DEN). For our analysis, we selected DEN-exposed tumors (32,494 substitutions).

#### Coordinates of genomic elements

The genomic coordinates of human protein coding genes (n=20,098) were obtained from ENCODE (http://www.gencodegenes.org) using GENCODE release v.19 (ftp://ftp.ebi.ac.uk/pub/databases/gencode/Gencode_human/release_19/gencode.v19.annotation.gtf.gz). Only coding sequences (CDS) of protein coding transcripts (‘gene_type’ and ‘transcript_type’ annotated as ‘protein-coding’) were retrieved as previously described in Mularoni *et al*., 2016. Promoter regions (n = 20,096) were obtained by mapping the 2500 bp sequences immediately upstream the transcription start sites (TSS) of protein-coding genes, and we removed all nucleotides that corresponded to protein-coding sequences, untranslated regions (UTRs) or short intronic splice sites (Mularoni *et al.*, 2016). C3H mice genomic coordinates for protein coding genes (n=19,234) version C3H_HeJ_v1 were downloaded from Ensembl release 90 (ftp://ftp.ensembl.org/pub/release-90/gtf/mus_musculus_c3hhej/Mus_musculus_c3hhej.C3H_HeJ_v1.86.chr.gtf.gz). Only CDS of protein coding transcripts (‘feature’ and ‘gene_biotype’ annotated as ‘CDS’ and ‘protein_coding’, respectively) were retrieved. Overlapping elements of the same type (CDS or promoters of the same gene) were merged together. Genomic elements showing incorrect annotations were manually removed from the datasets.

#### Implementation

OncodriveCLUSTL is implemented as a Python 3.5 package. It depends on external Python libraries including *bgreference*, *click*, *daiquiri*, *intervaltree*, *matplotlib*, *numpy*, *pandas*, *scipy*, *statsmodels* and *tqdm*. OncodriveCLUSTL takes as input a TSV file containing mutational data of multiple samples of either WXS or WGS data and a TSV annotations file containing the genomic positions of the genes or other genomic elements to be analyzed. OncodriveCLUSTL generates two main output files: 'elements_results.txt' contains a list of genomic elements ranked by the significance of their clustering signal; 'clusters_results.tsv' lists the information of all clusters found in genomic elements deemed significant. For a detailed explanation on how to run OncodriveCLUSTL please check the README document at https://bitbucket.org/bbglab/oncodriveclustl.

#### Cancer Gene Census list

We obtained the latest version of the COSMIC Cancer Gene Census (CGC) list containing a total of 719 genes (https://cancer.sanger.ac.uk/census) (downloaded on October 15$^{th}$, 2018) (Sondka *et al.*, 2018). Among them, 574 and 145 genes were classified in COSMIC Tier 1 and Tier 2, respectively. Tier 1 genes are those genes with solid available evidence of activity relevant to cancer as well as evidence of mutations in cancer that promote oncogenic transformation; Tier 2 genes correspond to genes with strong indications of a role in cancer but with less extensive available evidence.

#### OncodriveCLUSTL methodological details

OncodriveCLUSTL analyzes clustering signals in nucleotide sequences from human and non-human data. It has been tested using the hg19 reference human genome, c3h, mm10, cast and car mouse genomes and f344 rat genome obtained via the *bgreference* package (https://bitbucket.org/bgframework/bgreference). In addition, OncodriveCLUSTL can virtually be run using data from any reference genome provided that it is compiled through the *bgreference* package (details in README).

**Input data parsing.** By default, OncodriveCLUSTL only requires two main inputs: (i) mutations file, a TSV file containing the substitutions identified across a cancer whole exome (WXS) or whole genome (WGS) sequencing cohort; (ii) annotations file: a TSV file with the genomic positions annotations of the genes or other genomic elements (GEs) to be analyzed (details in README). By default, OncodriveCLUSTL analyzes all GEs provided in the annotations file. Alternatively, OncodriveCLUSTL can limit the analysis to a number of user-specified GEs (details in README). To start the analysis, OncodriveCLUSTL first parses the GEs coordinates from the annotations file. GE mutations are obtained by intersecting its genomic coordinates with those of substitutions (SNVs).

**Nucleotide context mutational probabilities calculation**. The computations of the significance of the clusters of mutations detected by OncodriveCLUSTL relies on a background model calculated from the input cohort nucleotide k-mer context mutational probabilities. By default, mutational probabilities of each k-mer based substitution are computed from the mutations file, under the assumption that all samples come from the same cancer cohort. Alternatively, OncodriveCLUSTL can calculate one specific mutational profile for each cancer type or cohort contained in the mutations file (e.g., Pancancer file). In addition, OncodriveCLUSTL is prepared to run using a mutational profile provided by the user (details in README). This option can be of interest when analyzing a cohort with few mutations.

To compute the mutational profile, OncodriveCLUSTL takes into account SNVs whose reference k-mer sequence does not contain a sequence gap or unannotated nucleotides in the reference genome. OncodriveCLUSTL can compute trinucleotide or pentanucleotide mutational probabilities. Briefly, a dictionary is constructed including all possible reference k-mers and 3 possible alternates of each of them (192 for trinucleotides, 3072 for pentanucleotides). Following the methodology laid out in Mularoni *et al*. (2016), mutational probabilities of each reference k-mer to each alternate k-mer are computed as the number of SNVs observed for the reference-alternate pair out of the total number of SNVs analyzed in the cohort or cancer type (Mularoni *et al*., 2016).

**Clustering analysis**. The following analysis was restricted to those GEs in the annotations file that contained more than 2 SNVs mapping the mutations file. We proceeded in the following steps:

1) ***Analysis of observed mutations***. OncodriveCLUSTL conducts spatial clustering analysis of the mutations observed along the GE. Two alternative modalities are available: i) by concatenating the connected components of the GE supplied (e.g., glueing the consecutive ends of the CDS chunks in the exons of a given gene) in which case we increase the method's sensitivity towards clusters in expanding the boundaries of the collapsed regions or ii) analyzing the connected components of the GE separately (by default). For both modes, the following subsequent steps are carried out:

(1a) ***Smoothing***. We used a Tukey kernel to smooth the function *n* that maps each position of the GE to the number of mutations observed at that position. Remark that the smoothing is carried out in the position coordinates implied by the chosen modality. Bearing that in mind, the smoothing function *S* is defined for each position *p* as:

$$S(p)=\frac{1}{M}\sum_{i=-L}^{L} n(p+i)T(\frac{i}{L});$$

where $T$ is the Tukey function defined as:

$$T(x) = (max\{1 - x^2, 0\})^2;$$

$M$ is the total mass spread by the kernel:

$$M = \sum_{i=-L}^{L} T(\frac{i}{L}) \sim L \int_{-1}^{1} T(x) dx;$$

and $L$ is the half window length where the smoothing is applied. The smoothing window length is 11 bp long by default.

(1b) *Root clusters generation*. Upon smoothing, root clusters of the GE are computed using the function *S,* which is defined along the GE's sequence. A cluster is always defined as 3 positions x-y-z in the linear sequence. For each local maxima **m** of S in the linear sequence we will define its root cluster as follows: i) if m is neither the first nor last position of the GE, then y=m, and x and z are as the closest local minima surrounding a local maxima y of *S;* ii) if m is the first positon of the GE, then x=y=m and z is the first local minima; iii) if m is the last position of the GE, then y=z=m and x is the last local minima.

(1c) *Merging of clusters*. In this step, the algorithm recursively identifies clusters that are closer than a given gap length (clustering window) and merges them in one new unified cluster. Starting from the 5'-most cluster in the GE defined as x-y-z, the algorithm looks for 3' adjacent clusters x'-y'-z'. The clusters merge if the distance between z and y' is smaller than or equal to the clustering window length. Then, a new cluster is obtained by merging both root clusters. The 5' and 3' boundaries of the new cluster are x and z', respectively. The maximum of the new cluster corresponds to the maximum with highest smoothing score amongst the root clusters, y or y'. When y and y' have equal scores, if they are contiguous positions y is selected as the new clusters maximum; otherwise, the clusters are not merged. The search iterates until no further merging is possible. The clustering window length is 11 bp long by default.

(1d) *Scoring of clusters*. A score is assigned to each cluster based on the number of SNVs it contains and their distribution across the cluster as follows:

$$Score = N \cdot \sum_{i} f_i \cdot 2^{-d_i/2};$$

where $N$ is the total number of SNVs mapping to the GE; $i$ runs through all mutated positions in the cluster; $f_i = 100 \cdot (m_i / N)$ is the percentage of mutations observed at position $i$ with respect to $N$; and $d_i = |i - i_{max}|$ is the distance from $i$ to the position reaching the maximum value of the smoothing function in the cluster. Given two clusters with the same number of mutations, this formula favours the one with the mutations concentrated in fewer positions. Clusters with fewer mutations than the defined threshold (2 by default) are scored to 0.

(1e) *Scoring of the GE*. The clustering score of a GE corresponds to the sum of the scores of its clusters.

2) ***Analysis of simulated mutations***. The same number of mutations observed in an GE are randomly sampled several times with replacement. The probability that a mutation is placed in a given position are derived from the mutational profile of the cohort, which is computed as explained above in the section *Nucleotide context mutational probabilities calculation*. Independently of the modality of observed mutations analysis (concatenated GE regions or not), mutations are randomized in the reference genomic sequence as follows. At the time of distribution of each mutation, a window of nucleotides (simulation window) centered at the mutated nucleotide is defined (default). The simulated mutation is therefore distributed at any position within the simulation window. To notice, when a mutation is observed in the region's boundaries, the simulation window centered at the mutated nucleotide may expand outside the region analyzed. Thus, simulated mutations can be sampled from outside the GE region. These simulated mutations, however, do not contribute to the simulated clusters and GE's scores, which ultimately affects the significance of the observed clustering signals. In addition, expanding simulation windows may not account for different mutational processes acting on regions of different nature (e.g., exons and introns, Frigola *et al*., 2017). As an alternative, if specified, OncodriveCLUSTL allows to displace the simulation window to fit inside the analyzed region in those cases where a subset of bp of the simulated window fall outside the analyzed region (e.g., if a window extends part of an exon and an intron, it can be placed to fit inside the exon). Displaced simulation windows maintain the defined window length but do not respect the central position for the observed mutation. For those cases where the length of the simulation window is greater than the region analyzed, the simulation window is trimmed to the region start and end. In this simulation mode, simulation windows that do not expand regions boundaries maintain the observed mutation in the central position. In our analysis, all simulations were done restricting simulations to the region studied. Once all mutations are randomly distributed across their respective simulation windows, OncodriveCLUSTL infers the clustering (steps 1a-1e) for the randomly generated mutations using the same analysis mode as in observed mutations (concatenated GE or not). Each iteration of random distribution of mutations (1,000 in all analyses described) retrieves simulated clusters and GE's scores.

**P-value computation and multiple test correction**. OncodriveCLUSTL generates three p-values per GE. First, an empirical p-value is computed as the fraction of iterations that yield a simulated GE score greater to or equal than the observed GE score. Second, an analytical p-value is calculated by fitting simulated GE scores to a gaussian kernel density estimate distribution and deriving the upper quantile of the observed GE score. Third, a second analytical p-value corresponding to the top-scoring cluster of the GE is computed following the same approach, fitting the distribution of the simulated cluster scores. To reduce the burden of analytical p-values computations, the algorithm randomly samples a subset of 1,000 simulated GEs over the total simulated GEs scores if the number of simulations is greater than 1,000; likewise, 1,000 simulated cluster scores are randomly sampled when the number of simulated clusters exceeds 1,000. All resulting p-values are subsequently adjusted (q-values) using the Benjamini-Hochberg method at 1% false-discovery rate (FDR). In other words, GEs with q-value ≤ 0.01 are considered potential drivers. All results shown here are based on rankings of GE scores analytical p-values.

#### OncodriveCLUSTL model selection

**Window hyperparameters: smoothing, clustering and sampling.** OncodriveCLUSTL is an unsupervised method to identify clustering of mutations along the genomic sequence of GEs. The method should raise a clustering signal whenever the clustering observed departs from what it would be expected assuming that the mutations are generated under neutral evolution. However, the method resorts to three main hyperparameters that strongly determine its performace: the shape of identified clusters depends on the smoothing (i) and clustering (ii) windows; the simulation of mutations depends on a sampling or simulation window (iii) which defines the region where mutations are randomly distributed. On the one hand, large smoothing and clustering windows tend to generate large clusters for either observed and simulated mutations; on the other hand, large simulation windows tend to spread simulated mutations along the sequence, which decreases the likelihood of formation of simulated clusters. The interplay between smoothing, clustering and sampling windows determines what kind of clustering signals the unsupervised method is bound to identify. Changes to the distribution of GEs p-values may end up affecting their goodness of fit to the uniform distribution.

We devised a strategy for model selection based on two criteria: i) goodness of fit of observed p-values vs. the uniform distribution; ii) enrichment of *bona-fide* known cancer elements in the ranking given by the method as an output. For each dataset, we ran OncodriveCLUSTL over all possible combinations in a predefined grid of hyperparameter values and selected the best configurations according to these criteria.

***Goodness-of-fit.*** For each configuration, we want to test whether the distribution of observed p-values is similar to the theoretical distribution of p-values, i.e., the uniform distribution in the interval [0, 1]. To this end, we computed the Kolmogorov-Smirnov (KS) goodness-of-fit statistic with respect to the uniform distribution in the interval [0, 1]. The KS statistic is defined in terms of two cumulative probability functions: i) the one arising from the observed data (empirical cumulative probability, ECP):

$$ECP(x) = \frac{1}{n}\sum_{i=1}^{n} 1_{(-\alpha, x]}(x_i);$$

And ii) the one arising from the uniform distribution (theoretical cumulative probability, TCP), which is linear. In order to proceed: first, we take the subset of observed p-values which are greater than 0.1; second, we randomly sampled 1,000 of them to avoid sample size biases in our comparisons; third, we calculated the KS statistic, which essentially measures the size of the maximum gap (deviance) between the observed and theoretical cumulative probability functions. To distinguish between p-value deflation and inflation, we computed whether the number of p-values above a threshold $\alpha = 0.1$ was greater (inflation) or lower (deflation) than expected. Thus we defined a signed version of the KS statistic which is positive for the inflated (resp. negative for the deflated). We selected all configurations bearing an the absolute value of the KS statistic up to 10% larger than the minimum KS statistic. This procedure left us with a set of most suitable configurations.

***Enrichment in bona-fide known cancer elements.*** For each configuration we calculated a CGC enrichment score for the top ranking genes (n=40). For each $1 \leq n \leq 40$, we computed the proportion of CGC genes (C for short) within the subset of genes with rank *n* or lower, hereinafter $S_n$. Then, we added up all these proportions, albeit giving more weight to the terms arising from smaller sets. Hence, we computed the following enrichment score:

$$E = \sum_{n=1}^{40} \frac{1}{\log_2(n+1)} \cdot \frac{|S_n \cap C|}{|S_n|}$$

The configuration with highest E was selected. For cohorts where this model selection strategy could not provide accurate models, we explored addtional values combinations of the before mentioned hyperparameters and manually curated the model selection on case-by-case basis using the approach described above. We hypothesize that the differences in performance could result from differential mechanisms shaping the mutational landscape for each cancer type (Alexandrov *et al.*, 2013). Therefore, we advice users to carry out a model selection according to their data specificities and constraints. Information of all the adjusted hyperparameters for all analyzed TCGA cohorts can be found in Supp. Table 1. The functions used to generate these data are available at https://bitbucket.org/bbglab/oncodriveclustl. For those cohorts where the CGC enrichment could not be applied (e.g., promoter regions, mouse data), model selection was based on the goodness-of-fit (Supp. Table 1).

We illustrate the performance of the KS test with the following example. We calculated the empirical cumulative distribution of p-values obtained by different configurations of the TCGA WXS UCEC cohort, using smothing windows of 11, 15, 21, 25, 31, 35, 41 and 45 bp; clustering windows of 11, 15, 21, 25 and 31 bp; and simulation windows of 31 and 35 bp. We randomly sampled n=100 p-values and calculated the ECPs between 0.1 and 1 for each of the configurations (Supp. Fig. 1). Finally, the best configuration was choosen as the one with higher CGC enrichment, as explained.

**Background mutational probabilities**. To test which of the k-mer context mutational probabilities, trinucleotides or pentanucleotides, generated more accurate models, we ran OncodriveCLUSTL using 3-mer and 5-mer contexts with the 19 selected TCGA WXS datasets, keeping the rest of parameters as default. We calculated the Kolmogorov-Smirnov (KS) statistic to assess the fitness of the observed distribution of p-values to the expected uniform distribution (Supp. Fig. 2). For these data, we found no clear differences between the KS statistic obtained from trinucleotide or pentanucleotide based background models. Given that trinucleotide mutational probabilities are less computationally expensive, we set them as the default. However, recently Martincorena and colleagues have shown that pentanucleotide contexts can explain more accurately the mutational processes in melanomas caused by UV light (Martincorena *et al.*, 2017). Consequently, all analyses shown in the main paper were carried out using the default trinucleotide mutational probabilities, except for the melanomas cohort, to which the pentanucleotide mutational probabilities was applied to compute the background model. We recommend users to conscientiously select OncodriveCLUSTL tri- or penta-nucleotide probabilities that best fit their own mutational datasets.

**Quantile-quantile plots.** In order to evaluate the models generated by OncodriveCLUSTL, we generated quantile-quantile plots (QQ-plots) comparing the observed p-value distribution (y-axis) with the expected uniform p-value distribution (x-axis). We consider a model to be accurate if the observed p-values closely fit the uniform distribution (red dash line) for the most part of the GEs analyzed, i.e., those GEs not bearing a significant clustering signal (red dots, $q < 0.01$). The names of the 10 top-ranking genes are shown. The names of gene symbols annotated in the CGC appear in bold. The code used to generate QQ-plots is available at https://bitbucket.org/bbglab/oncodriveclustl.

#### Running examples

Complete guideliness on how to run OncodriveCLUSTL can be found in the README doument at https://bitbucket.org/bbglab/oncodriveclustl. Briefly, we highlight here different running options of the algorithm that can be selected through the command line:

*Default run:*
~$ oncodriveclustl -i /INPUT_PATH/.../mutations_file.tsv -o /OUTPUT_PATH/output_directory -r /INPUT_PATH/.../regions_file.tsv.gz

*Human coding sequences using default parameters:*
~$ oncodriveclustl -i /INPUT_PATH/.../mutations_file.tsv -o /OUTPUT_PATH/output_directory -

r /INPUT_PATH/.../regions_file.tsv.gz --concatenate

*Human coding sequences using non-default parameters:*
~$ oncodriveclustl -i /INPUT_PATH/.../mutations_file.tsv -o /OUTPUT_PATH/output_directory -r /INPUT_PATH/.../regions_file.tsv.gz --concatenate -simw 35 -sw 35 -cw 15 -kmer 5

*C3H mice coding sequences using default parameters:*
~$ oncodriveclustl -i /INPUT_PATH/.../mutations_file.tsv -o /OUTPUT_PATH/output_directory -r /INPUT_PATH/.../regions_file.tsv.gz --concatenate -g c3h

#### Benchmark

The performance of OncodriveCLUSTL in coding regions was assessed by computing the enrichment of the genes identified by the method for the set of CGC genes with at least 2 SNVs in the cohort under analysis. To compare it with OncodriveCLUST (Tamborero *et al.*, 2013) and HotMAPS (Tokheim *et al.*, 2016), we calculated the fold increase in the proportion of CGC genes among sets with increasing number of top ranking genes. Briefly, for each set of increasing number of top ranking genes, we calculated the fraction between the proportion of GCG genes within the set and the proportion of CGC genes within all genes detected by the method. This correction accounts for the fact that methods differ in the number of genes analyzed and therefore the proportion of analyzed GCG genes among them, which ultimately modifies the probabilities of detecting CGC genes. Enrichment plots show the enrichment of the top 40 ranking genes identified by OncodriveCLUSTL, OncodriveCLUSTL and HotMAPS.

To show the complementarity between linear (2D) and non-linear (3D) clustering methods we calculated the number of significant CGC genes ($q < 0.01$) detected by OncodriveCLUSTL and HotMAPS. For each cohort, we computed the number of unique CGC genes detected by OncodriveCLUSTL, and HotMAPS, as well as the number CGC genes detected by both. In parallel, we tested the complementarity of 2D linear clustering to other methods based on different signals of positive selection, including OncodriveFML (Mularoni *et al*., 2016) for functional impact bias and dNdScv (Martincorena *et al*., 2017) for recurrence. To this end, OncodriveCLUST version 1.0, HotMAPS version 1.1.3, OncodriveFML version 2.1.0 and dNdScv version 0.1.0 were run using default parameters.

#### Cluster plots

We generated the named “cluster plots” to illustrate the distribution of mutations, smoothing curve and clusters along the sequence of a GE. The plot shows GEs sequence 5’-3’ (left to right) in the strand encoding them. For those GEs fragmented in different regions (e.g., exons in a gene), the sequence is shown as concatenated (x-axis). The different regions are delimitated by the different white-grey chunks and dashed lines. The first grid of the plot contains the number of mutations (left y-axis) mapped to their relative position in the concatenated GE. Smoothing curves (blue) correspond to the smoothed values per position following Tukey's kernel density estimate application (right y-axis, labels not shown). GE score and p-value are highlighted in a box. The second grid illustrates observed clusters. Significant clusters are highlighted in a red-color scale where darker red corresponds to more significant p-values. Non-significant clusters are shown in grey. All functions used to generate cluster plots are included in OncodriveCLUSTL code and can be run through the command line (details in README).

#### Expression analysis

Expression and copy-number data from TCGA WGS-505 dataset were obtained from Fredriksson *et al.* 2014. As described in the original article, RNA-sequencing (RNA-seq) BAM format data (hg19 assembly) and copy-number amplitudes (Affymetric SNP6 platform) of CDS (n=20,167) and lncRNAs (n=11,852) (GENCODE v17; Harrow *et al*., 2012) were processed following the methodology introduced by Akrami *et al*., 2013. Pancancer differential expression analysis between non-mutated and cluster-mutated samples was carried out for samples bearing a diploid copy number of the gene under analysis (log2 absolute amplitude < 0.2). Differences were assessed using U-Mann Whitney test ( $\alpha=0.05$ ).

#### Code availability

OncodriveCLUSTL algorithm and a running example are available under GNU Affero General Public License at https://bitbucket.org/bbglab/oncodriveclustl. OncodriveCLUSTL version 1.0 was used to generate the results shown in the manuscript and supplementary material. Further versions will be available through our repository. Additionally, a limited version of OncodriveCLUSTL can be run through our web at https://bbglab.irbbarcelona.org/oncodriveclustl.

### 2 Supplementary Figures

**Supp. Fig. 1.**

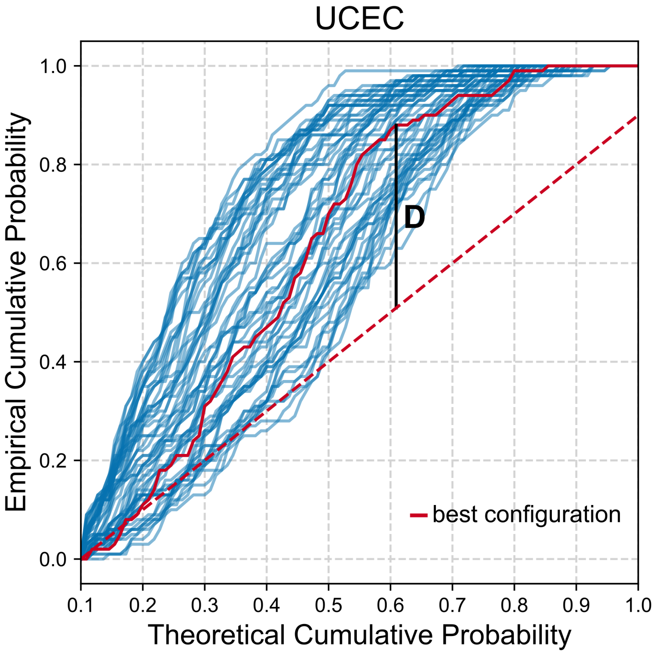

**Supp. Fig. 1.** Representation of the empirical cumulative probabilities of p-values for TCGA WXS UCEC cohort. For each configuration, the empirical cumulative probability of p-values was calculated and plotted against the theoretical uniform cumulative distribution between 0.1 and 1. The striped red line shows the expected uniform cumulative distribution. Results for the best configuration of parameters given the top 40 CGC enrichment are highlighted in red. The vertical black line shows the maximum deviance (D) of the best configuration. An alpha of 0.05 was used.

**Supp. Fig. 2.**

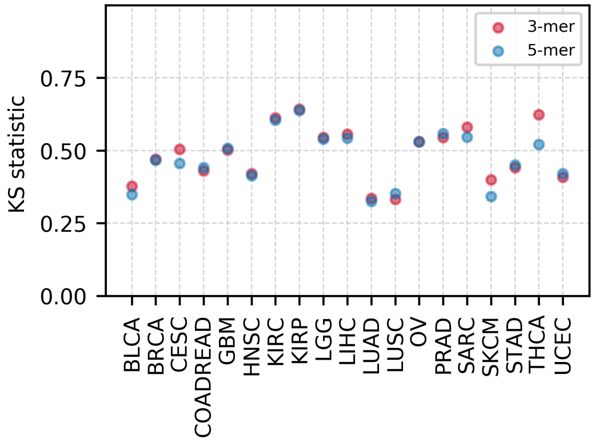

**Supp. Fig. 2.** Effect of trinucleotide or pentanucleotide context mutational probabilities on OncodriveCLUSTL models. The Kolmogorov-Smirnov statistic showing the fitness of the observed p-values distribution to the expected uniform distribution was calculated for the 19 TCGA cohorts analyzed trinucleotide (3-mer, red) and pentanucleotide (5-mer, blue) context mutational probabilities. Smoothing and clustering parameters were ran as default.

**Supp. Fig. 3.**

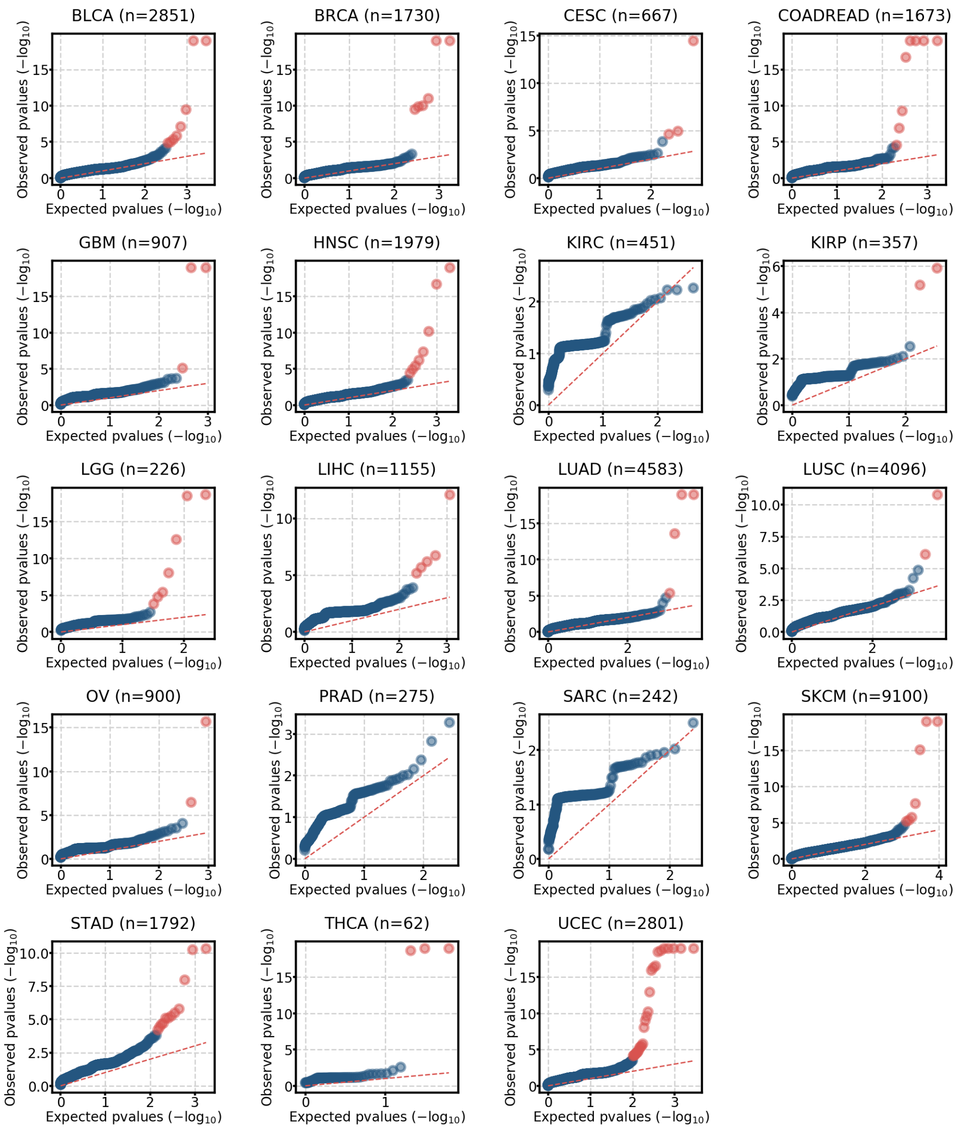

**Supp. Fig. 3.** Model adjustement by OncodriveCLUSTL. QQ-plots showing the observed p-values distribution versus the expected uniform distribution for all TCGA cohorts analyzed. Genes with q < 0.01 are highlighted in red. The number of p-values plotted on each QQ-plot is shown on top.

**Supp. Fig. 4.**

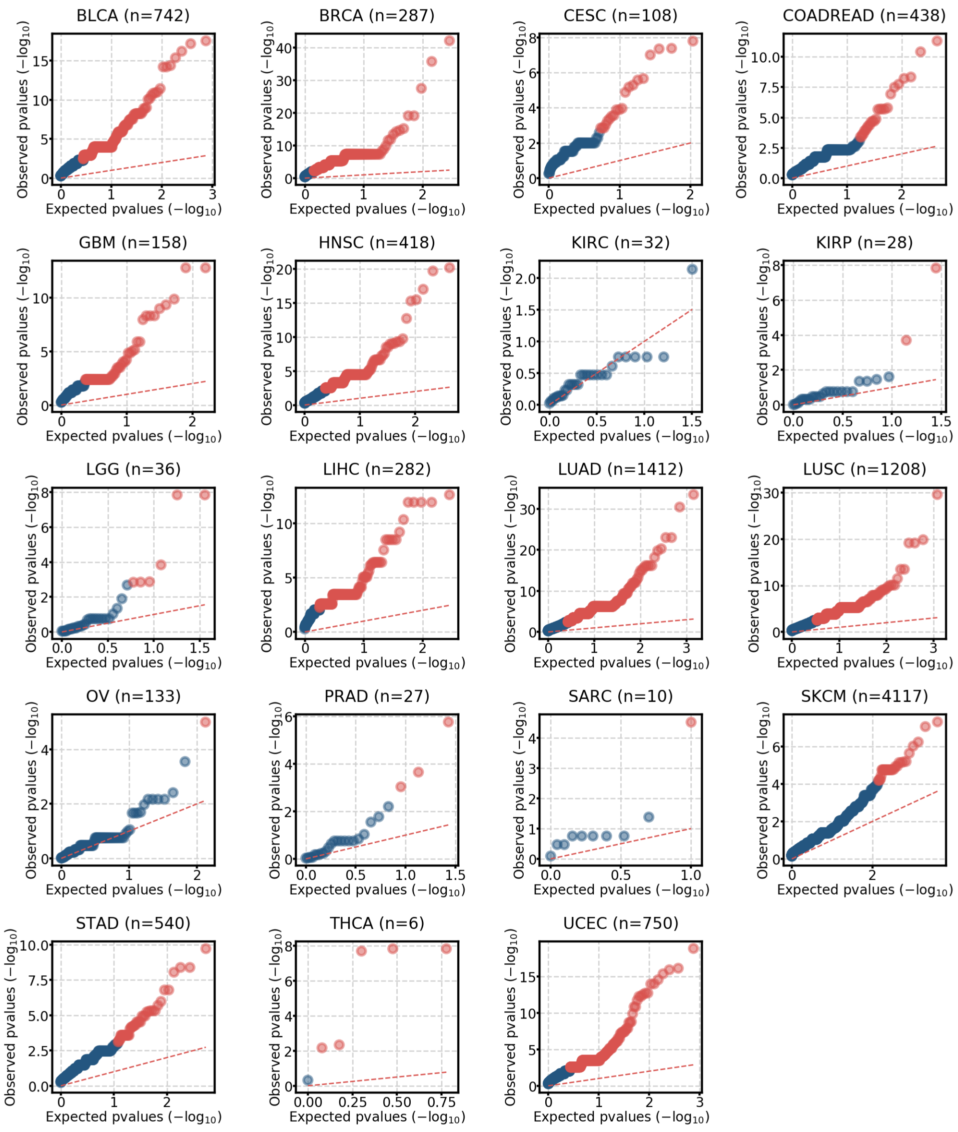

**Supp. Fig. 4.** Model adjustement by OncodriveCLUST. QQ-plots showing the observed p-values distribution versus the expected uniform distribution for all TCGA cohorts analyzed. Genes with q < 0.01 are highlighted in red. The number of p-values plotted on each QQ-plot is shown on top.

**Supp. Fig. 5.**

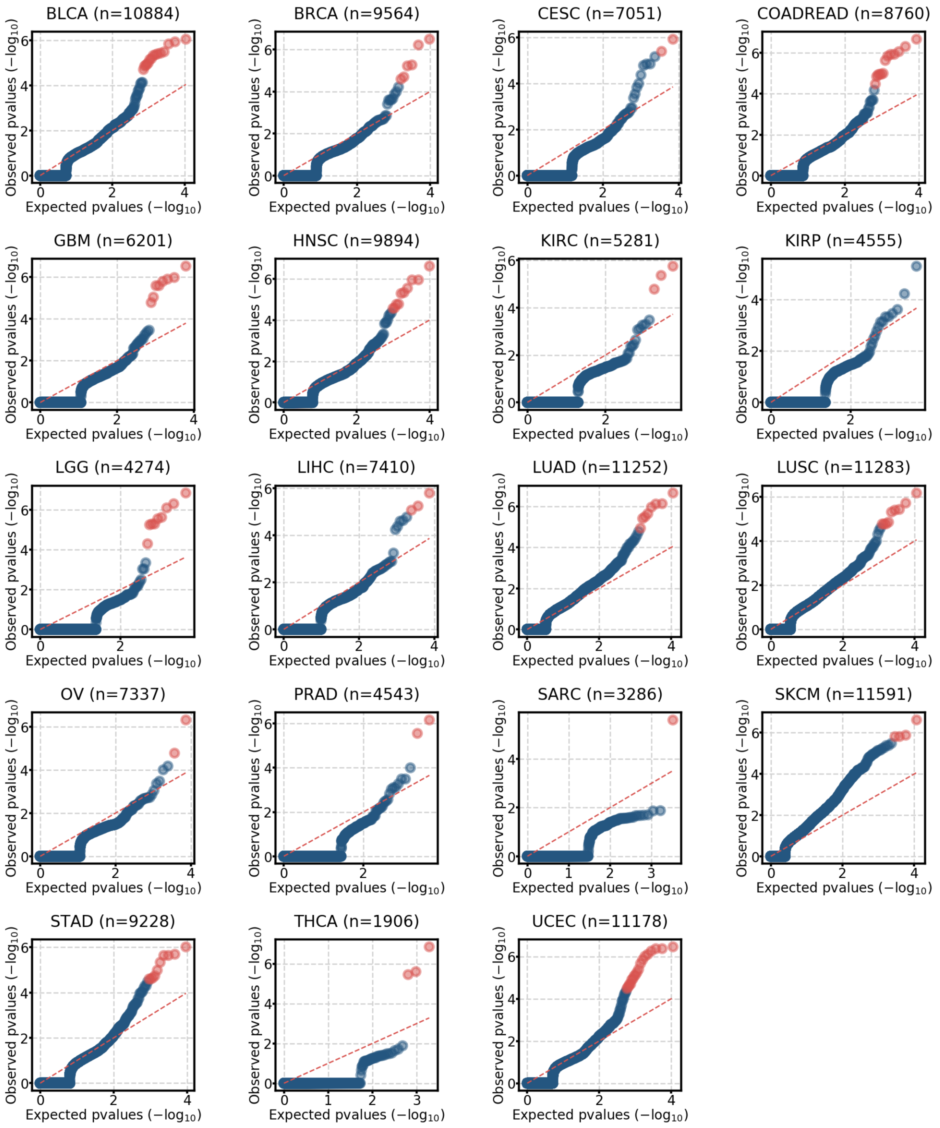

**Supp. Fig. 5.** Model adjustement by HotMAPS. QQ-plots showing the observed p-values distribution versus the expected uniform distribution for all TCGA cohorts analyzed. Genes with q < 0.01 are highlighted in red. The number of p-values plotted on each QQ-plot is shown on top.

**Supp. Fig. 6.**

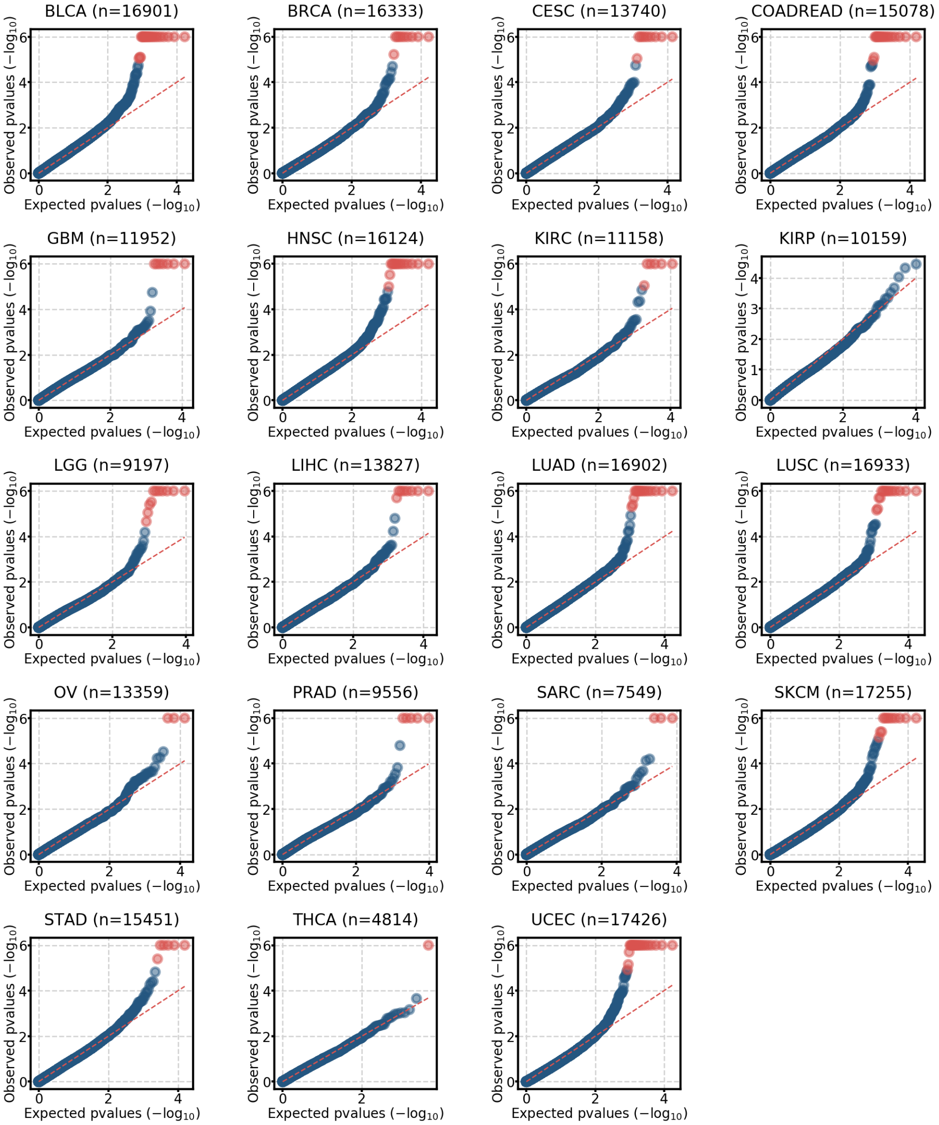

**Supp. Fig. 6.** Model adjustement by OncodriveFML. QQ-plots showing the observed p-values distribution versus the expected uniform distribution for all TCGA cohorts analyzed. Genes with q < 0.01 are highlighted in red. The number of p-values plotted on each QQ-plot is shown on top.

**Supp. Fig. 7.**

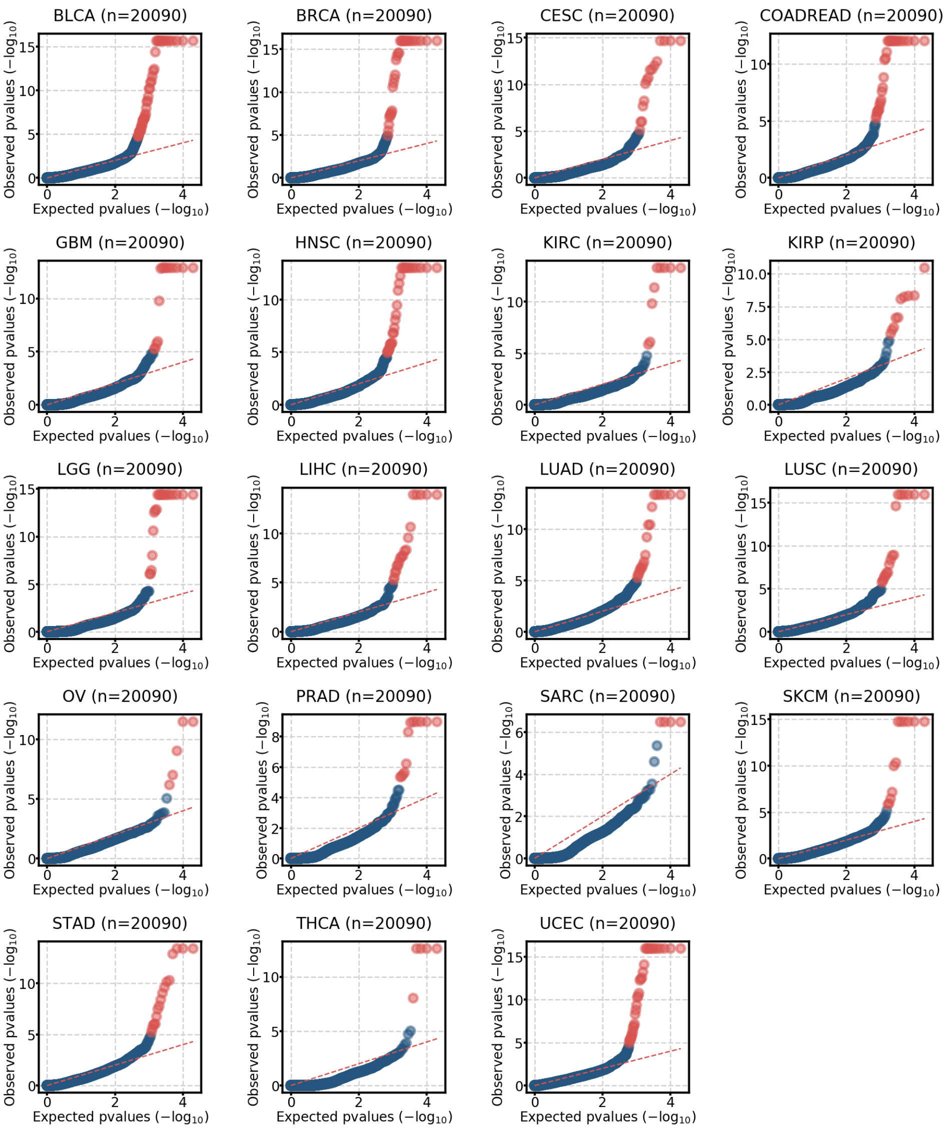

**Supp. Fig. 7.** Model adjustement by dNdScv. QQ-plots showing the observed p-values distribution versus the expected uniform distribution for all TCGA cohorts analyzed. Genes with q < 0.01 are highlighted in red. The number of p-values plotted on each QQ-plot is shown on top.

**Supp. Fig. 8.**

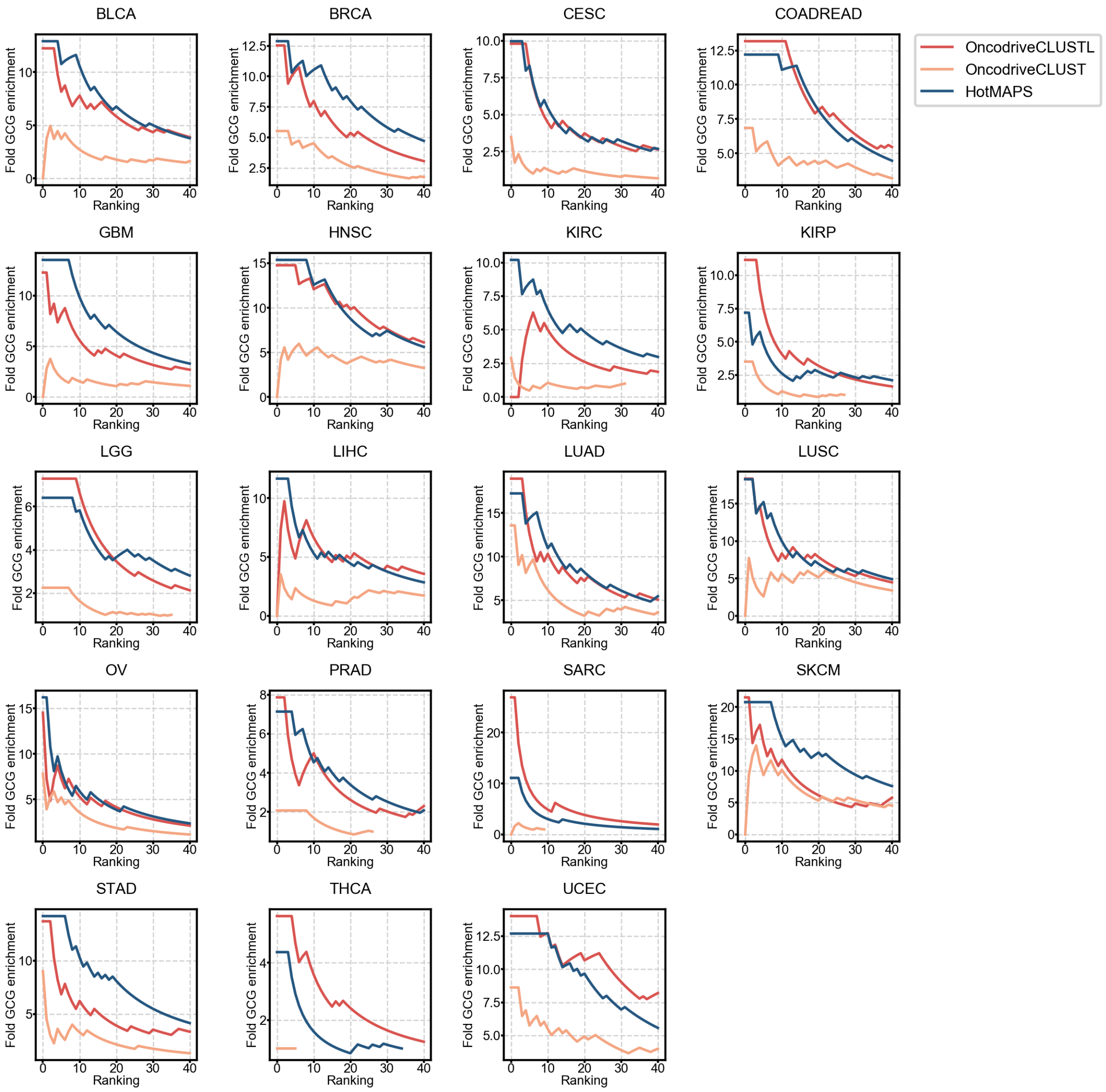

**Supp. Fig. 8.** CGC genes enrichment of OncodriveCLUSTL, OncodriveCLUST and HotMAPS. The fold increase in the proportion of CGC genes among sets with increasing number of the top 40 ranking genes was calculated for OncodriveCLUSTL, OncodriveCLUSTL and HotMAPS.

**Supp. Fig. 9.**

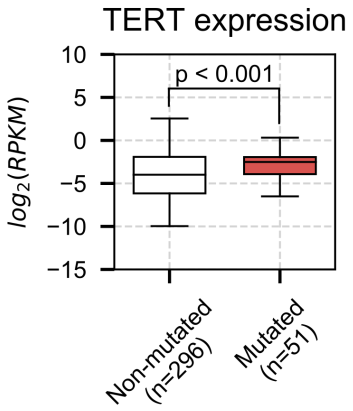

**Supp. Fig. 9.** TERT expression analysis. Samples containing a mutation in the chr5:1295228-1295253 cluster (red) detected by OncodriveCLUSTL have an increased TERT expression when compared to TERT promoter non-mutated samples (U-Mann Whitney $p < 0.001$). Only those samples with non-altered TERT gene copy number alterations were used for this analysis.

### 3 Supplementary Tables

**Supp. Table 1.** OncodriveCLUSTL running parameters for the tested cohorts. Best smoothing, clustering and simulation windows (bp) OncodriveCLUSTL according to the model selection strategy introduced in Supplementary Methods section.

### 4 Supplementary references

Akrami,R. *et al.* (2013) Comprehensive Analysis of Long Non-Coding RNAs in Ovarian Cancer Reveals Global Patterns and Targeted DNA Amplification. *PLoS One*, 8, e80306.

Alexandrov,L.B. *et al.* (2013) Signatures of mutational processes in human cancer. *Nature*, 500, 415–421.

Connor,F. *et al.* (2018) Mutational landscape of a chemically-induced mouse model of liver cancer. *J. Hepatol.*, 69, 840–850.

Ellrott,K. *et al.* (2018) Scalable Open Science Approach for Mutation Calling of Tumor Exomes Using Multiple Genomic Pipelines. *Cell Syst.*, 6, 271–281.e7.

Fredriksson,N.J. *et al.* (2014) Systematic analysis of noncoding somatic mutations and gene expression alterations across 14 tumor types. *Nat. Genet.*, 46, 1258–1263.

Frigola,J. *et al.* (2017) Reduced mutation rate in exons due to differential mismatch repair. *Nat. Genet.*, 49, 1684–1692.

Harrow,J. *et al.* (2012) GENCODE: the reference human genome annotation for The ENCODE Project. *Genome Res.*, 22, 1760–74.

Martincorena,I. *et al.* (2017) Universal Patterns of Selection in Cancer and Somatic Tissues. *Cell*, 171, 1029–1041.e21.

Mularoni,L. *et al.* (2016) OncodriveFML: A general framework to identify coding and non-coding regions with cancer driver mutations. *Genome Biol.*, 17, 128.

Sabarinathan,R. *et al.* (2016) Nucleotide excision repair is impaired by binding of transcription factors to DNA. *Nature*, 532, 264–267.

Sondka,Z. *et al.* (2018) The COSMIC Cancer Gene Census: describing genetic dysfunction across all human cancers. *Nat. Rev. Cancer*, 18, 696-705.

Tamborero,D. *et al.* (2013) OncodriveCLUST: Exploiting the positional clustering of somatic mutations to identify cancer genes. *Bioinformatics*, 29, 2238–2244.

Tokheim,C. *et al.* (2016) Exome-scale discovery of hotspot mutation regions in human cancer using 3D protein structure. *Cancer Res.*, 76, 3719–3731.

Zou,X. *et al.* (2017) Short inverted repeats contribute to localized mutability in human somatic cells. *Nucleic Acids Res.*, 45, 11213–11221.
